# Evolution of colonial life history in styelids tunicates involves changes in complexity patterns

**DOI:** 10.1101/2020.05.31.126409

**Authors:** Stefania Gutierrez

## Abstract

Biological complexity is defined as the number of modules that compose an organism or a biological system, the type of interactions between these modules, and new hierarchies that describe these interactions. These patterns in biological complexity are changing during the evolution of life-histories, such as the evolution of coloniality in animals. In relation to coloniality, it is possible to observe an increment in all the aspects defined in the concept of biological complexity. First, in colonial animals, there is an increment in the modules that compound the system (i.e. zooids) compared with a solitary organism in which the multicellular individual a unity. Consequently, this transformation of the multicellular individual, in a component of the modular architecture in colonies, involves an increase in the regulatory processes of colonial system. This is precisely the case of the colonial life history evolution from solitary ancestors in the Styelids tunicates. Therefore, the main question of this study is How is the regulation of the asexual developmental processes that occurred simultaneously in the modules of the colonies? This question was studied, by the research of colonial strategy in the styelid *Symplegma.* Using in vivo observations of the budding process, description and classification of the extra-corporeal blood vessels system and the blood cells, by cytohistological assays. The conclusion is that the regulation of the simultaneous developmental processes that occurred in *Symplegma* colonies is mediated by the system of extra-corporeal blood vessels, which maintain physically the cohesion of the individuals, the plasma, and migratory blood cells transport signals between the individuals of the colonies.

## 1. Introduction

> “Evolutionary change involves the increasing complexity of a feature already present in ancestors”
>
> — Stephen Jay Gould, from *Ontogeny and Phylogeny,* 1977

Gould (1977, pp 268-269) suggested that an increase in complexity is an inevitable condition for evolution by acceleration of developmental processes. Complexity in biology is used to refers number of cell types, body parts, or biological processes (McShea 1996)□. Increase or reduction in number of these characters, results in changes of complexity patterns (McShea 2017)□.

Complexity is a useful concept to understand how developmental mechanisms, environmental conditions and natural selection are acting in evolution of new life histories. Clarifying, this does not imply a directionality in evolution. On the contrary, the idea is that complexity is a useful concept to understand the evolution of new life histories- “We will be interested in the whole pattern of change, not just the increases … but also the decreases, the frequent retreats into simplicity”-as McShea (2017, pp 2) proposed. In biological systems complexity is represented by the number of modules (e.g. cell type, leg-pair type, zooids, polyps) that compose an organism, the type of interactions between these modules, new biological hierarchies or nestedness processes that describe these interactions (McShea 1996, 2017; Adami 2002)□, and also the capacity of self-organization in biological systems (Yaeger 2009).

Evolution of colonial life history in animals is one example of the change in the complexity patterns. These changes are observed in the modularization of multicellular individuals, forming the colony as a new biological hierarchy (Davidson et al, 2004)□; or by new types of interactions of these modules, such as the cellular migration and the molecules involved in communication between modules (Lauzon et al., 2007)□; or by self-organization, such as the rearrange of extracorporeal blood vessels to maintain homeostasis after a disturbance (Rodriguez et al. 2017)□.

### 1.2. Evolution of colonial life history in Styelidae from a solitary ancestor, implies the increase of characters complexity in colonial descendants

Tunicates are useful to understand changes in patterns of complexity related to the colonial life history. Colonial tunicates evolved multiple times, in pelagic and benthic environments (Kocot et al., 2018).□ Such as, in the tunicate family Styelidae, colonial life history evolved by convergence two times from solitary ancestors (Alié et al. 2018)□. Evolution of colonial animals involved the increase in the number of modules, interactions, hierarchical processes and self-organization of the colonies (Fig.1) (Ballarin et al. 2008; Gutierrez and Brown 2017; Alié et al. 2018)□. Although, evolution of colonial life history increased the complexity patterns in Styelids.

**Figure.1.**
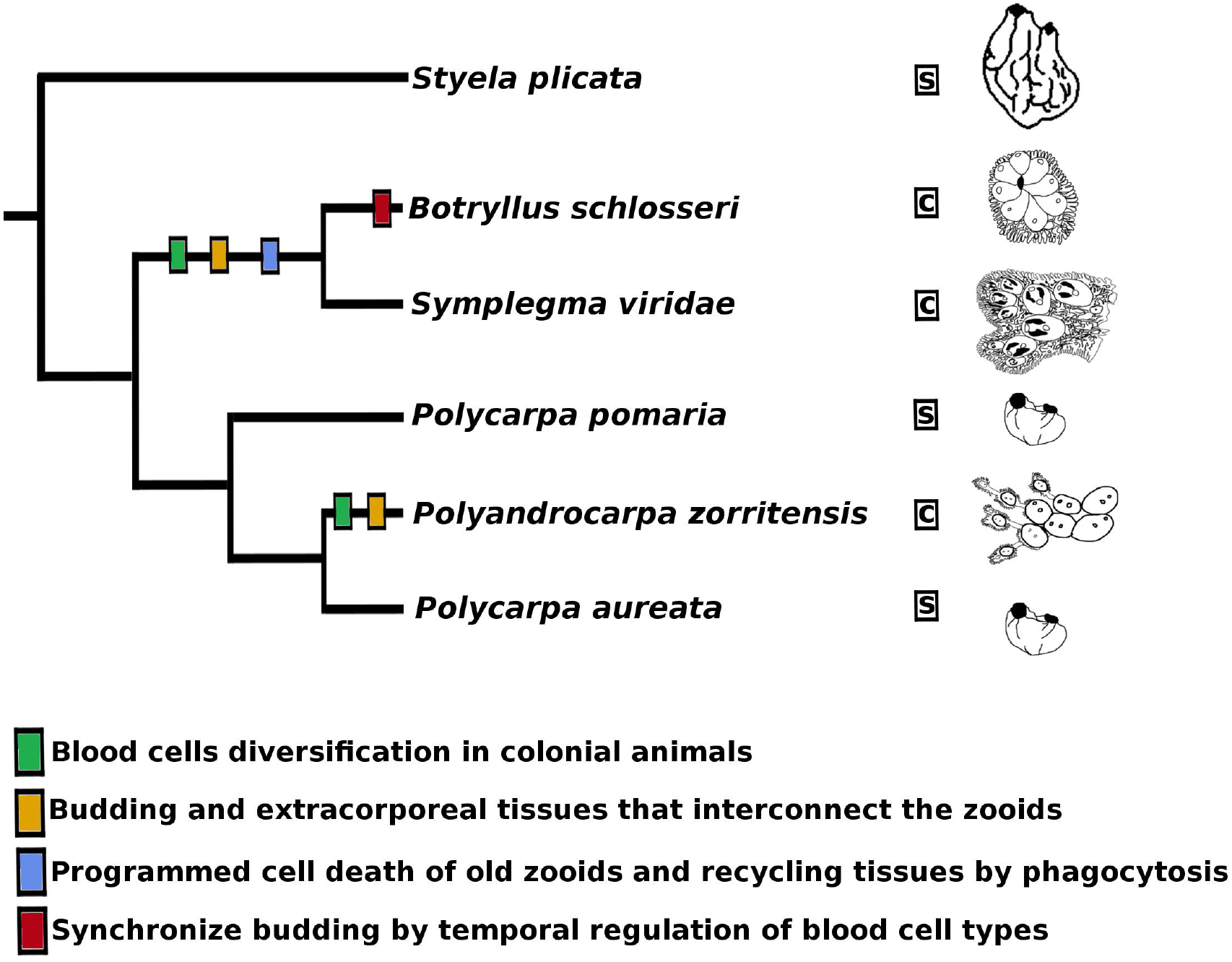
Phylogenetic summary of the characters related with colonial life history evolution in Styelidae. The proposed key characters for evolution of coloniality are: the specialization of blood cells; blood cells diversification to manage modules in colonies; budding and extra-corporeal tissues to interconnect zooids; programmed cells death of old zooids and their tissues recycling; the synchronized budding in botryllids. Based in Alié et al., 2018; Hiebert et al., 2019 and Zeng et al., 2006

Genus *Symplegma* is one of the colonial animals in Styelidae. *Symplegma* colonies are composed by zooids, and a extra-corporeal vessel system with specialized blood cells that circulate constantly(Mukai & Taneda, 1978)□. New buds appear from the evagination of vessels and epitheliums from adult zooids. Blood vessel system is very dynamic, buds move during development. Buds formed from adult zooids, move far away from parental zooid, by the formation of a new vessel to connect to the general vessels system of the colony. Vessels have the capacity to rearrange in case of different disturbance and external stimulus. Such as, in the whole-body regeneration, blood vessels pump blood by themselves, allowing cells circulation and regeneration (Sugimoto and Nakauchi 1974; Gutierrez and Brown 2017)□. In absence of young buds, by systematical removals, new buds are formed and developed faster. This suggest that the asexual development of zooids is modulate by mechanisms at the colony level (Gutierrez and Brown 2017)□.

Solitary life form is consider the ancestral character in Styelidae (Kott 2005; Zeng et al. 2006; Alié et al. 2018)□. Thus, evolution of *Symplegma* colonial strategy, involved the develop of more complex characters in comparison with solitary forms. Modularization of multicellular individuals in colonies, implies modularization of developmental processes (e.g morphogenesis and aging). These developmental processes are occurring simultaneously, one of the main innovation of the colonies in comparison with the solitary forms (Jackson & Coates, 1986; Jackson & Hughes, 1985). These processes are spatially and temporary impossible in solitary animals. Evolution of colonial life history is related with the increase of complexity in characters, such as extra-corporeal vessels; diversification of blood cells; and regulatory mechanisms to modules coordination.

The developmental processes are modularized in *Symplegma* zooids. Simultaneously a bud is forming, by a process analogous to a grastrulation; another bud is differentiated its organs; the fully differentiated zooid is filtering and feeding the colony; and an old zooid is aging and dying (Kawamura and Nakauchi 1986; Gutierrez and Brown 2017)□. The main question of this study is try to understand How is the regulation of the asexual developmental processes that occurred simultaneously in *Symplegma* colonies?

The premises to answer this question are: (a) all the modules of the colony are interconnected by a blood vessel system in which blood cells are in constant circulation (b) there are specific type of blood cells related with main biological processes of the colonies, such as phagocytosis, budding and regeneration, allorecognition and storage cells, (c) external disturbances such as systematical remotion of modules (zooids or buds), cause changes in the proportion of the type blood cells, and alterations in the asexual development of the colony (Cima et al, 2001; Franchi et al., 2016; Gutierrez & Brown, 2017; Lauzon et al., 2007).□

Therefore, the proposed hypothesis is that t*he regulation of the simultaneous developmental processes that occurred in Symplegma colonies are mediated by the modularization of these developmental processes. To coordinate these simultaneous processes new blood cells types evolved diversifying in relation with biological functions associated to specific developmental stages (e.g progenitor cells related to early buds, phagocytes related to old zooids). In Symplegma clade evolved a cellular based communication system where signals are transmitted between modules (i.e zooids) by migratory blood cells and molecules diffused in plasma.*

To test this hypothesis the development of *Symplegma rubra* and *S.brakenhielmi* was observed, from the oozoid to adult colonies, to understand the formation of a colony from a first module. The colonial strategy in *S.rubra* was unknown, thus *S.rubra* budding and the blood cell types were characterized, comparing with the information reported before for *S.brakenhielmi.* Some blood cell types have an active behavior and fast movements. These cellular behaviors were observed and record in videos, to understand cellular behaviors associated with blood cell types. Blood cells were identified and counted, in the morphogenesis stages and aging stages. Comparing the types of blood cells at these these simultaneous stages. Finally the aging process of the zooids was describe to identify if is a programmed cell death, like in other colonies,or a senescent process.

## 2. Materials and Methods

### 2.1. Colonies and budding characterization

*Symplegma rubra* and *S. brakenhielmi* colonies were collected from floating structures in the Yatch Club IlhaBela-YCI (Ilhabela, São Paulo, Brazil). Fragments of colonies were attached to microscope slides and kept in open cages. Cages were immersed in the water from floating docks for three weeks. Grown colonies were cleaned and transported to the laboratory at Universidade de São Paulo. Colonies were maintained at 25° C and fed with a mixture of living algae (*Isochrysis, Thalassiosira, Pavlona, Nanochlorpsis)* and commercial food. Colonies were observed under stereomicroscope Leica M205 FA. Colonies at the reproductive stage were transported to the Centro de Biologia Marinha da Universidade de São Paulo – *CEBIMar.* The larvae released from the colonies were obtained and transferred to microscope slides to observe them.

### 2.2. Blood cell characterization

For the blood cells extraction was followed a previously described protocol (Cima 2010)□, with some changes to improved obtained results (See supplementary material 5.3 chapter 4). Colonies were immersed in anticoagulant for 5 minutes. Then ampullae were gently cut and blood was extracted. Anticoagulant was washed by centrifugation (10 minutes-3000 rpm), and the blood cells were resuspended in a solution of 1/3 anticoagulant-2/3 filtered sea water (FSW). The blood cells were attached to microscopy slides coated with Poly-L-Lysine. Afterwards the attached blood cells were stained using cytological techniques to observe the diversity and classify. Living blood cells were stained with a neutral red solution (8mg/l in FSW) to observe acid cellular compartments. Blood cells were fixed with 4% paraformaldehide in FSW and stained with Hematoxylin and eosin (H&E) or Giemsa 10%, to identify the cellular morphology. Blood cells were observed and photographed using the inverted microscope Leica DMi8.

### 2.3. Cellular behavior

Blood cells were collected as mentioned above. The cells were added to coversilps coated with laminin (50 μg/mL) and filmed under a microscope Zeiss AxioVert A1, equipped with the Canon DSLR camera. Images were recorded every 3 seconds and processed in the photo editing free software Darktable.

### 2.4. Blood cells identification in the bud morphogenesis and aging zooids

*Symplegma rubra* and *S. brakenhielmi* colonies were fixed in paraformaldehide 4% in FSW overnight and afterwards washed with PBS. Fixed colonies were dehydrated by ethanol series (25%, 50%, 70%, 80%, 90%, 100%) and two xylol washes and embedded in paraffin to be sectioned. Serial sections of 5μm thickness, perpendicular to the longitudinal axis of the zooids were obtained using a microtome. Sections were mounted on glass slides, deparaffinized and rehydrated with the inverse ethanol series mentioned before. The tissue slides were stained with H&E, mounted with Entellan and examined under a Zeiss AxioVert A1 light microscope. The proportion of the types of blood cells was calculated by counting the cells in the early stages of budding (double vesicle, stage.5) and in the aging stages of the old zooids (stage 11). These buds and zooids were in the same colony, thus the analyzed stages were present simultaneously in the colonies. All the blood cells inside the zooids and buds were counted in four sections every 20 μm, in the smallest buds were counted two sections every 10 μm. Tukey and Bonferrni tests were used to determine significant differences between blood cells proportion in the budding stages.

### 2.5. Characterization of aging zooids

Old zooids were observed *in vivo* in colonies of *Symplegma rubra and* colonies of *S. brakenhielmi.* From these *in vivo* observations and the resorption stages reported in *Botryllus schlosseri* were established homologous stages in *Symplegma*. The morphology of these resorption stages was described by histological slides stained with H&E as mentioned above.

## 3. Results

### 3.1 Development of a *Symplegma* colony

*Symplegma rubra* and *S.brakenhielmi* are species with a similar strategy of coloniality. The colonies are formed by modules (zooids and buds) interconnected by a systems of blood vessels with circulating blood cells. The formation of buds is a constant process and the budding occurs asynchronously in contrast with other species, such as the botryllids. *Symplegma* colonies are characterized by the formation of extension and growth zones. The location of the extended ampullae and the buds (Fig. 2A-B). Growth area redirects the growth of the colony. Colonies were capable of small movements and the positions of the zooids change dynamically across the blood vessels system.

**Figure.2.**
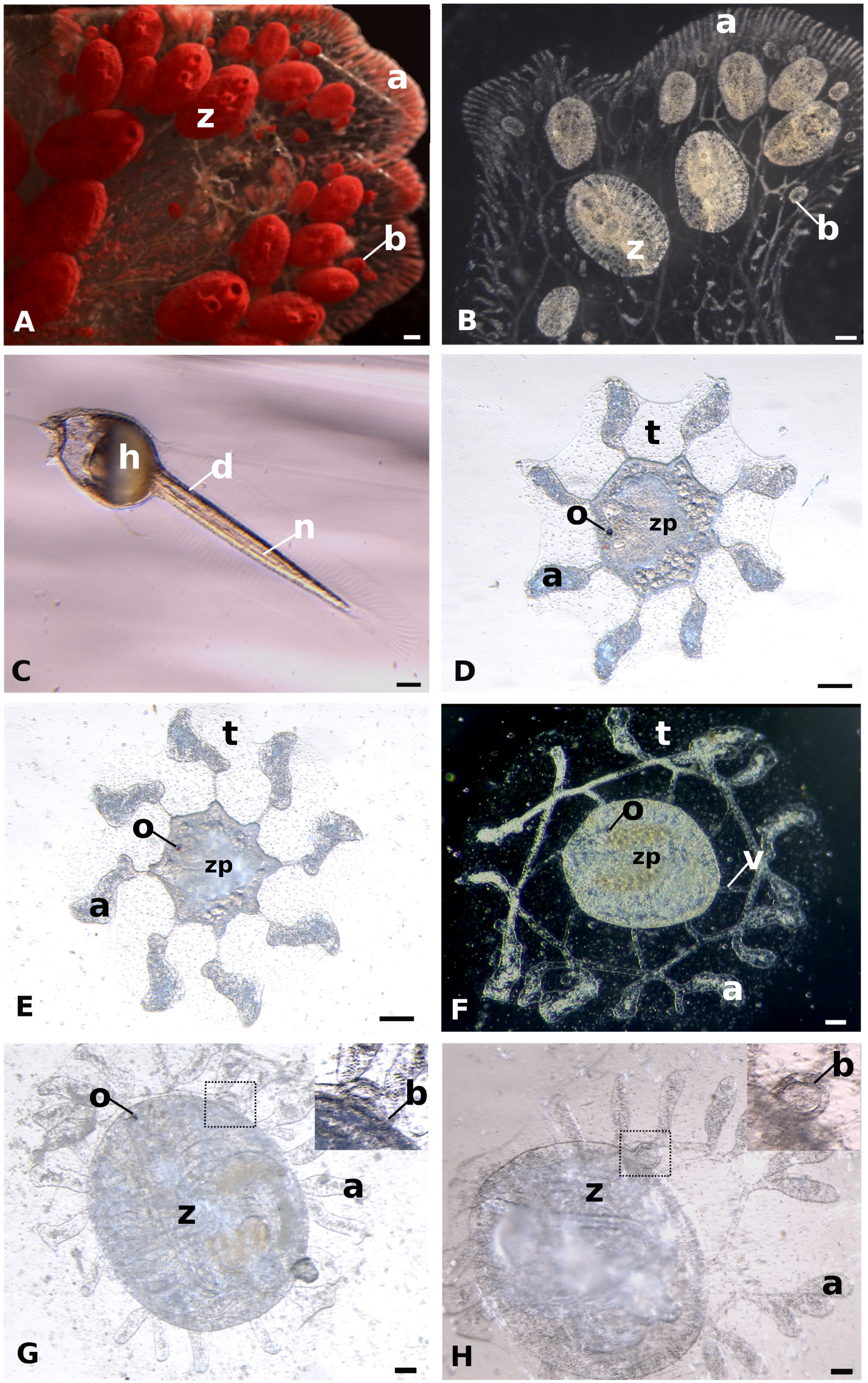
Coloniality in the genus *Symplegma*. (A) *Symplegma rubra* colony characterized by its red color. (B) Symplegma brakenhielmi colony characterize by its greenish color. (C) Larva of *S. rubra* after be released by the colony. (D) Larva of *S. rubra* after the settlement and metamorphosis (E) Larva of *S. brakenhielmi* after the settlement and metamorphosis. (F) Ozooid of *S. rubra* during the formation of the blood vessels syste*m.* (G) Formation of the new bud in *S. rubra.* (H) Formation of the new bud in *S. brakenhielmi. a: ampullae; b:bud; o: ocellus; t: tunic; v: blood vessel. Otholito; z: zooid; zp: zooid primordium.* Scale bar is 500 μm in A and B; 250 μm in C, D, E and F; 350 μm in G and F.

The colonies were reproductive during all the year, specially from December to February,the summer season at the south of Brazil. The reproduction is by internal fertilization and the larval development by brooding. Fully developed larva is approximately 2mm long, and has a circular head with sensorial papillae in the anterior part The tail has the notochord, the dorsal nerve and muscles. The beating of the tail propels the larva, which can swim for up to 12 hours before settlement (Fig. 2C). The metamorphosis starts with the resorption of the tail and the formation of the first zooid (i.e oozoid). At the beginning of the oozoid development some larval structures are remanent such as the ocellus and otolith (Fig. 2D). In *Symplegma brakenhielmi* and *S.rubra* the eight primordial ampullae form a remarkable symmetric pattern (Fig. 2D-C). Inside of these primordial ampullae, blood cells were circulating. Simultaneously with the formation of pharynx and internal organs of oozooid, the ampullae were growing and fusions forming the primordial blood vessels system (Fig. 2F). After a week of the settlement the blood vessels system ramifies and the fully differentiated oozoid open the siphons to feed. One day after, the siphon apertures were observed on the first buds (Fig. 2G -H). The lifespan of the oozoid is approximately 20 days, then the zooids and buds are developing and the blood vessels are forming. It is the beginning of the formation of a colony.

The formation of the colonies was very similar between the two *Symplegma* species, however the budding in *S.rubra* is by palleal budding (i.e buds are formed by the evagination of the pharyngeal and external epitheliums of the parental zooid). In contrast to *S.rubra,* in which budding is exclusively palleal, *S.brakenhielmi* has simultaneously palleal and vascular budding.

Budding in *Symplegma rubra* was characterized by eleven stages. The stages 1 to 3 are the formation of the budlet from the parental bud. Stage 4 is the beginning of folding of internal epitheliums to form the organs, analogous to grastrulation. Organogenesis occurs during stages 5 to 8, with tissues differentiation and formation of all the internal organs. Finally in stage 9 the zooid is fully functional and starts to filter. Lastly, the zooid starts a senescent process and it is resorb by the colony. The stages were established following Berrill (1941)□ and Sabbadin (1955)□ nomenclature and compare with the homologous stages previously reported in *S. brakenhielmi* and botryllids.

### 3.2. Blood cells of Symplegma rubra

The blood cells in *Symplegma rubra* are composed by a variety of cellular populations, characterized by cytological morphology. Some of the populations are cells with characteristics of precursors. The other populations were characterized in three functional lineages: phagocytes, cells for allorecognition, and storage cells (Fig. 3AB).

**Figure. 3.**
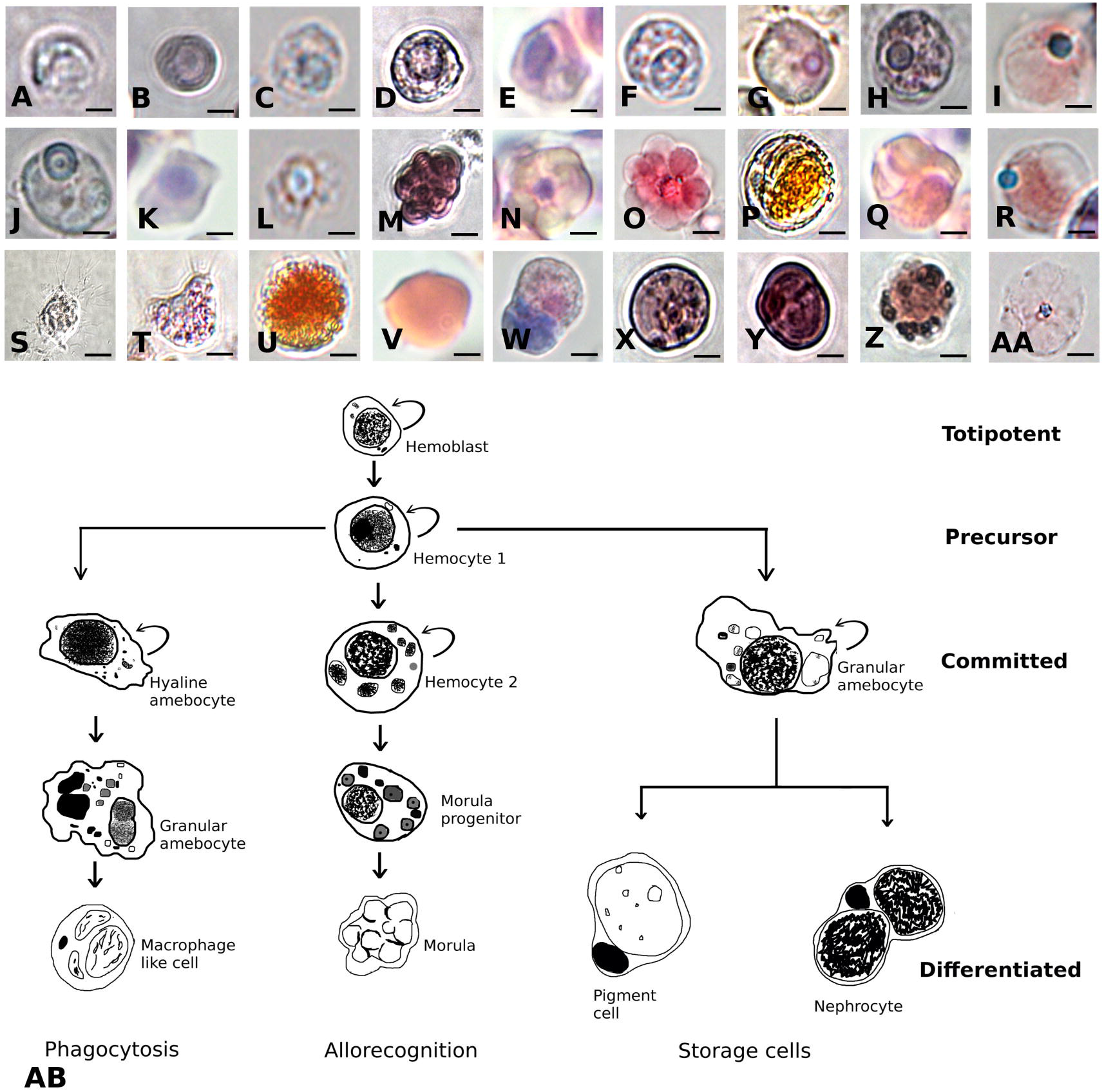
Hypothetical model of hematopoiesis in *Symplegma*. In *Symplegma rubra* were observed blood cell types similar to *Symplegma brakenhielmi* blood cells and another colonial tunicates. Living hemoblast stained with neutral red (A). Fixed hemoblast stained with Hematoxylin and eosine (H&E) (B) and giemsa (C). Living hemocyte 1 stained with neutral red (D). Fixed hemocyte 1 stained with H&E (E) and giemsa (F). Living hemocyte 2 stained with neutral red (G). Fixed hemocyte 1 stained with H&E (H) and giemsa (I). Living hyaline amebocyte stained with neutral red (J). Fixed hemocyte 1 stained with H&E (K) and giemsa (L). Living morula stained with neutral red (M). Fixed hemocyte 1 stained with H&E (N) and giemsa (O). Living macrophage-like cell stained with neutral red (P). Fixed macrophage-like cell stained with H&E (Q) and giemsa (R). Living granular amebocyte stained with neutral red (s). Fixed macrophage-like cell stained with H&E (T). Living nephrocyte stained with neutral red (U). Fixed nephrocyte stained with H&E (V) and giemsa (W). Living pigment cell stained with neutral red (X-Y). Fixed pigment cell stained with H&E (Z) and giemsa (AA). The hypothetical model of hematopoiesis is based in the cellular morphology of blood cells, frequency and proliferation observed in S. *brakenhielmi* and *S.rubra.* The hemoblast is proposed as the totipotent hematopoietic stem cell. The hemocyte 1 is propose as the precursor of the undifferentiated comamitted cells. These blood cells are the progenitors of the there genera blood cell lineages, phagocytes, allorecognition and storage cells. Scale bar is 1 μm in A and B; 2 μm in C and from X to AA; 3 μm from E to P, and from T to W; 4 μm in Q and R; 6 μm in S.

*Hemoblasts (HE):* cells with a size between 3-4 μm. HEs have a round regular cytoplasm with a high nucleus-cytoplasmic ratio. The cytoplasm has a small number of organelles, for that HEs are negative to neutral red (this dye stains cellular acid compartments). The nucleolus is clearly visible with hematoxylin and eosin (H&E) stain. The nucleus is stained blue with Giemsa (Fig. 3A-C).

*Hemocyte (H1):* cells with a size between 5-6μm. H1s have a regular cytoplasm with a high nucleus-cytoplasmic ratio and small number of organelles. The living H1s are negative to neutral red, and a strong basophilic stain in the nucleus with H&E. The nucleus and small vesicles are stained blue with Giemsa (Fig. 3D-F).

Hemocyte (H2): cells with a size between 6-7 μm. H2s are round shaped and have some granules in the cytoplasm. In living H2s the cytoplasm is stained with neutral red suggesting the content of acid compartments. The eccentric circular nucleus is characterized by a strong color in all the dyes (neutral red, H&E, and Giemsa) (Fig. 3G-I).

*Hyaline amebocyte (HA)*: cells with a size between 4-6 μm. HAs have a irregular amoeboid cytoplasm with small number of organelles. HAs are negative to neutral red dye an the cytoplasm has a clear color in all dyes. HAs have a high nucleus-cytoplasmic ratio, nucleus has a strong basophilic color with H&E and blue color with Giemsa (Fig. 3J-L).

*Morula cell (MC):* cells with a size between 8-10 μm. MCs have a irregular cytoplasm full of homogeneous round vesicles. MCs are positive to neutral red dye, suggesting an acid content in their vesicles. Cytoplam has a strong brownish-dark red color in all the dyes. The nucleus is small stained purple with H&E and blue with Giemsa (Fig. 3M-O). MCs have an active movement (Video. 1).

*Macrophage-like cell (MLC):* cells with a size between 10-12 μm. MLCs have huge vacuoles and some heterogeneous vesicles. MCs are positive to neutral red, suggesting acid content in the vacuoles. MCs have strong yellowish-orange color with H&E and Giemsa. The nucleus is eccentric and small, visible with Giemsa (Fig. 3P-R).

*Granular amebocyte (GA):* cells with a size between 10-15 μm. GAs have a irregular amoeboid cytoplasm with pseudopods. The cytoplasm contains granules with light colors stained with H&E and negative to neutral red (Fig. 3S-T). GAs have an active movement (Video. 2).

*Nephrocyte (N):* cells with a size between 8-20 μm. Ns have a round or hourglass shape. The cytoplasm contains dense granules with Brownian movement. Ns are stained brownish-yellowish with neutral red, H&E and Giemsa, suggesting acid content in the cytoplasm (Fig. 3U-W).

*Pigment cell (PC):* cells with a size between 8-15 μm. PCs have a irregular cytoplasm with granules inside. PCs in *Symplegma rubra* are red in color, probably related with the characteristic color of this specie. Some granules are stained with neutral red (Fig. 3X-AA).

This diversity of blood cells in Symplegma colonies, is associated with specific cellular behaviors. Morula cells and amebocytes show dinamic movements, specially amebocytes with the pseudopods. These faster cellular movement can be involved in the biological processes and communication between zooids in colonies (Video 1.-2).

### 3.3. Buds morphogenesis

In *Symplegma rubra* the right side of the peribranchial epithelium of buds is thick. From this thickening a budlet starts to form in a parental bud stage 5. This budlet expand and growth forming a double vesicle (stage. 3) from the peribranchial epithelium and external epithelium of the parental bud (Fig. 4A,C). The bud separates from its parental zooid and a new blood vessel is forming from the bud to the colonial systems of vessels (Fig. 2B). The bud increase in size and the internal vesicle starts to fold, forming the pharynx and stomach primordium (Fig. 4D). Buds in *S. brakenhielmi* are formed by palleal budding and from vessels as was reported before (Gutierrez & Brown, 2017)□. The palleal body in *S.brakenhielmi* and *S. rubra* follow the same pattern.

**Figure.4.**
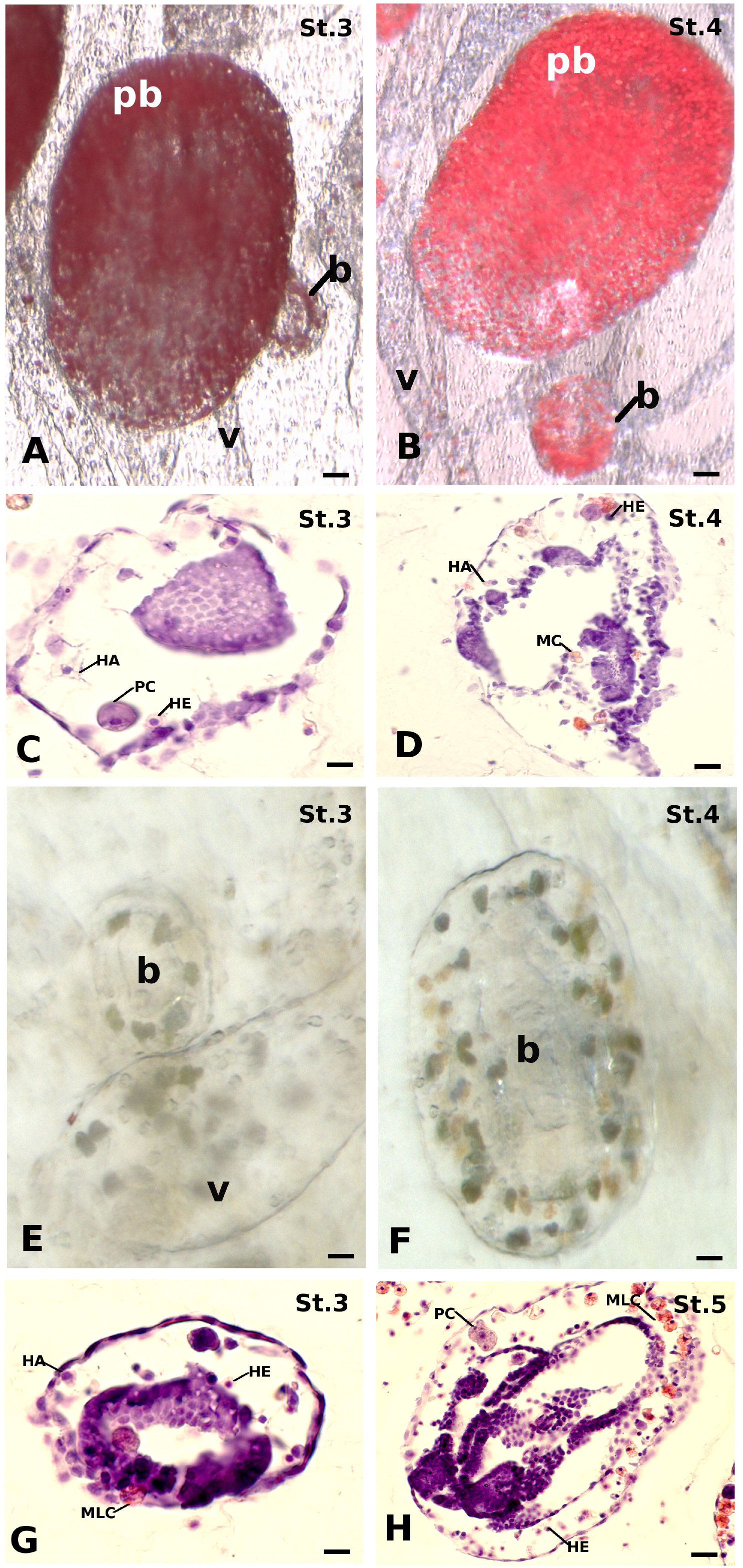
Buds morphogenesis in *Symplegma*. (A) The buds in *Symplegma rubra* are forming by paleal budding (i.e. buds are forming by the evagination of branchial and external epitheliums in the parental bud). Bud in the stage 3 or double vesicle is attached to its parental bud in stage 6. (B) The bud in stage 4 moved apart from the parental bud and it is attached to a near blood vessel. (C) Bud in stage 3 of *S. rubra,* the internal vesicle in in formation. Mostly of the cells observed inside the bud are hyaline amebocytes (HA), hemoblasts (HE) and a pigment cell (PC). (D) Bud in stage 4 of *S. rubra* HA, HE, MLC are between the external epithelium and the internal epithelium, at this stage there are morula cells (MC). (E) Buds in *Symplegma brakenhielmi* can be formed by paleal budding or vascular budding as is shown. (F) Vascular bud from a *S.brakenhielmi* colony. (G) Bud in stage 3 of *S. brakenhielmi.* Stage known as double vesicle has inside HA, PC and HE, also Macophage-like cells are entering in the bud. (H) Bud in stage 3 of *Symplegma brakenhielmi.* Stage known as double vesicle has inside HA, PC and HE, also Macophage-like cells are entering in the bud. *b:bud; pb: parental bud; v: blood vessel.* Scale bar is 50 μm in A and B; 10 μm in C; 20 μm from D to H.

Blood circulation is observable inside buds since double vesicle stage (St.3) and continues during all the asexual development. Blood cells with characteristics of precursors (i.e as hemoblasts and hyaline amebocytes) were observed in the morphogenesis of buds. During the bud development increase the number of macrophage-like cells (Fig. 4C,D,G,H). Hemoblasts, hyaline amebocytes and pigment cells were statistically more frequent in double vesicles than in old zooids for the two *Symplegma* species (Fig. 6).

### 3.4. Resorption of old zooids

Stage 11 is the final stage in *Symplegma* budding. The lifespan of the fully differentiated zooid is between 3 and 3.5 days in the two *Symplegma* species. The first step of the final stage 11.1 is the closure of siphons. When touched siphons did not react, as mentioned by Ballarin et al., (2008) for *Botryllus schlosseri* (Fig. 5A, D). After twelve hours the stage 11.2 started, with the longitudinal antero-posterior contraction of the body. (Fig. 5 B, E). During this stage pharynx and stomach epithelia initiated a disintegration, by the separation of epithelial cells. Macrophage-like cells began to accumulate around pharynx and stomach (Fig. 5G). At stage 11.3 the size of the zooid reduced dramatically, and the heart beating was slower. The disintegration of the internal organs was clearly observable, and the body was inundate by macrophage-like cells. The blood that circulate around the old zooids was denser than around young zooids. These macrophage-like cells were seen moving outside from the old zooid, probably with the resorbed tissues. Finally mostly of the tissues of the old zooid were phagocyted and moved outside from the body and the heart beating stopped (Fig. 5C, F, H). Remanent of tunic and tissues stayed during two days before the resorption was complete. During the resorption stage the proportion of macrophage-like cells was significantly bigger than in young bud of the same colony. In *Symplegma rubra* the proportion of morulas was significantly bigger than in young bud (Fig. 6). The resorption process has similar pattern in the two *Symplegma* species.

**Figure.5.**
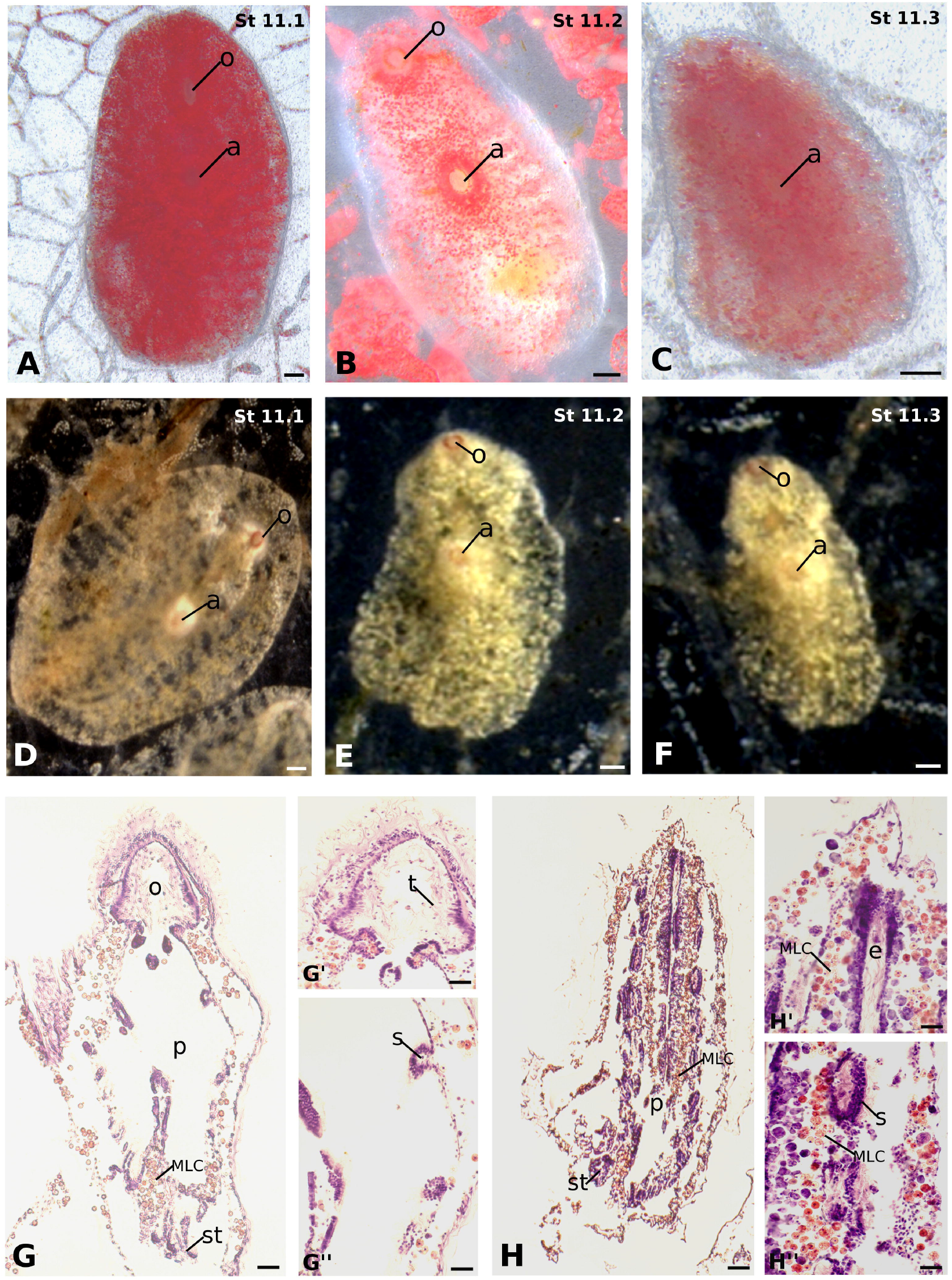
Resorption of old zooids in *Symplegma*. The resorption process of old zooids is similar in th species *Symplegma rubra* and *Symplegma brakenhielmi*. (A,D) Old zooid at the stage 11.1 (12 hours), the first step of resorption. Oral siphons closure and starting the antero-posterior contraction. (B,E) Old zooid at stage 11.2 (12 hours). The contraction of body continues and the zooid reduces size. It is possible to observe an empty tunicate, probably because many cell including pigment cells were phagocyted and moving outside from the old zooid. (C,F) Stage 11.3 the zooid continue reducing its size, the tunicate is more transparent and the hear beat is slower and stops progressively. (G) Old zooid starting the anterior-posterior contraction, the oral siphon is already close. Macrophage-like cells (MLC) are ingression and accumulation around the pharynx and the stomach. (G’) Magnification of the closed oral siphon. (G’’) Magnification of the pharynx, which start by the disintegration process of the epitheliums. (H) Old zooid in an advance resorption process. The body contraction is increasing. MLCs are increasing their proportion around the pharynx and the stomach. (H’) Magnification of the endostile. (H’’) Magnification of the stigmas. *a: atrial siphon; e: endostyle; o: oral siphon; MLC; macriphage-like cell; p: pharynx; s: stigma; st: stomach; t: tentacles.* Scale bar is 250 μm from A to D; 600 μm from E to F; 50 μm G and H; 30 μm in G’ and G’’; 20 μm in H’ and H’’.

**Figure.6.**
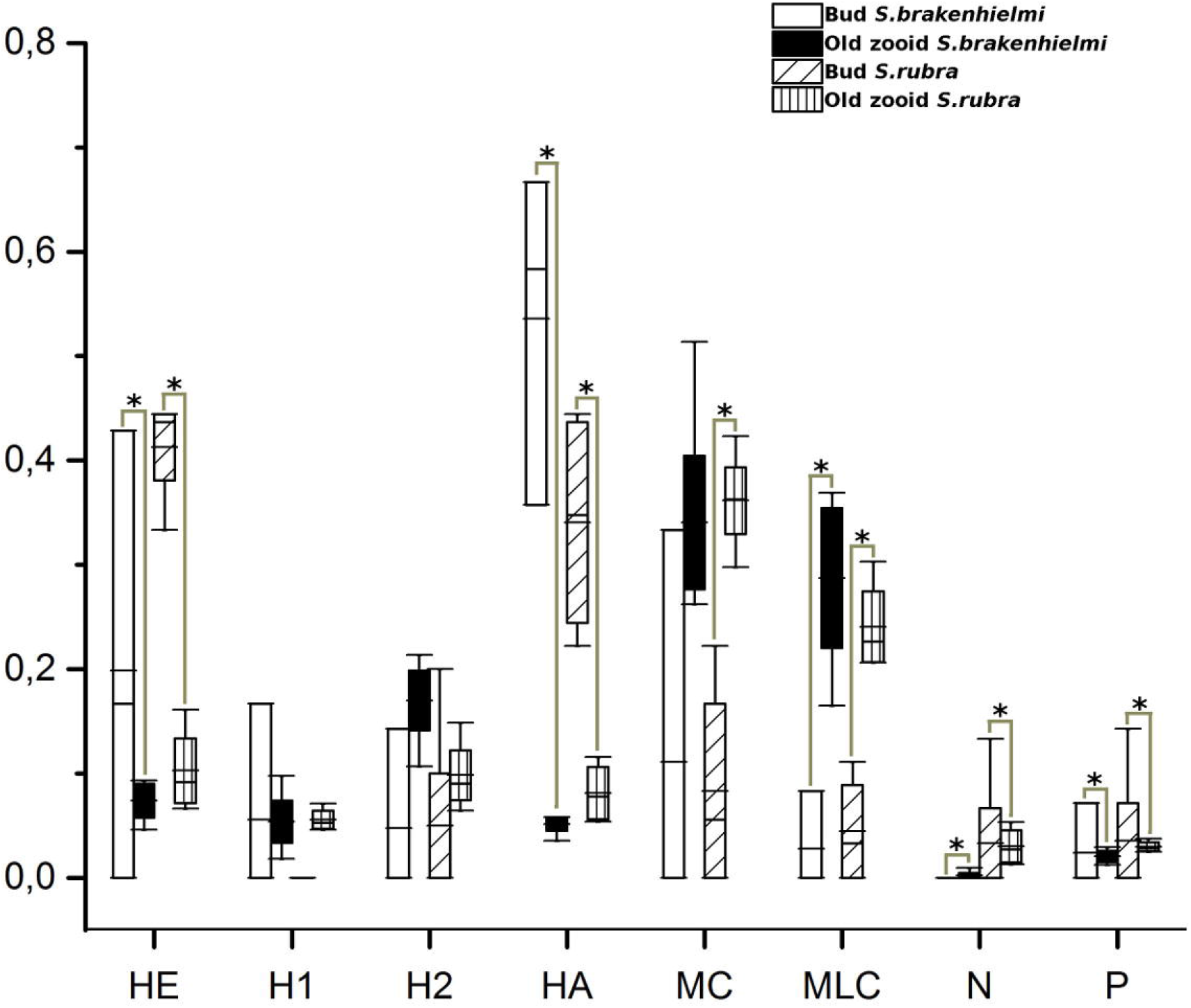
Proportion of blood cells in buds and in aging zooids. The proportion of blood cells is different between buds and aging zooids. Blood cells with undifferentiated characteristic as HE and HA are statistically more frequent in buds than in aging zooids. In contrast in aging zooids are more frequent MLC and MC. The proportion of storage cells (N,P) is variable. Pigment cells are statistically more frequent in buds, moreover nephrocytes have different results between *Symplegma* species. *S. brakenhielmi* has more Ns in aging zooids than in buds, the contrary to *S.rubra.*

**Figure.7.**
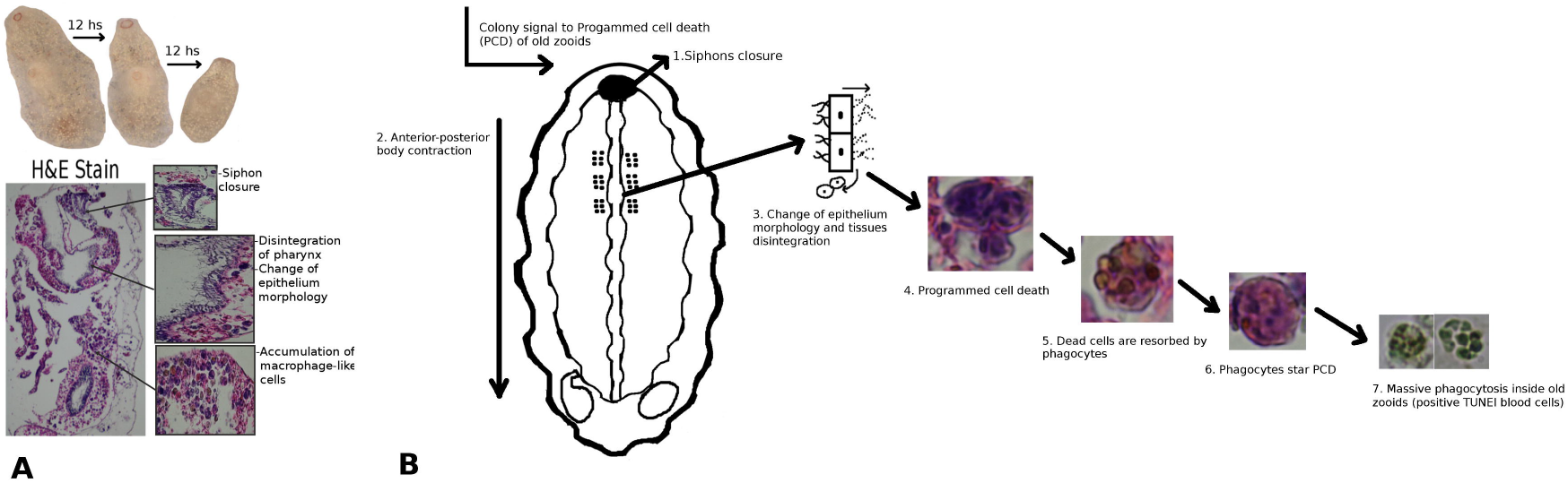
Hypothetical model of aging in *Symplegma*. (A) The aging in *Symplegma* colonies begins with the siphon closure followed by the anterior-posterior contraction around the longitudinal axis of the zooid. Macrophage-like cells (MLC) migrate to the zooid and the internal tissues start to disintegrate. (B) After the epithelial disintegration, these cells start apoptosis. The apoptotic bodies are phagocyted by the MLCs that migrated and increase the proportion MLCs in the aging zooid. Some types of blood cells show signals of DNA fragmentation by TUNEL, inside the aging zooid and the vessels. The hear-beating decrease and finally the internal tissues are phagocyted, and probably this phagocyted material is recycled by the colony.

## 4. Discussion

### 4.1. Modularity of a multicellular individual by the development of a colony of zooids

The colonies in *Symplegma* have defined areas, characterized by specific biological processes: extension area, with the younger buds in morphogenesis; area of fully differentiated zooids; and the regression area, with the aging zooids (Fig. 2A-B). The extension area is more evident in colonies that are growing, like the cultured colonies that were growing from their natural substratum to the glass slides. This suggests that the extension area redirect growth of the colony, probably related with the perception of good environmental conditions received for ampullae. The oozoid in development and the primordial ampullae form a symmetric pattern with eight ampullae. This remarkable pattern was conserved in *S.rubra* and *S.brakenhielmi* oozoids (Fig. 2 D-E).

Simultaneously with the development of the oozoid, beginning the formation of blood vessels system by the fusion of primordial ampullae (Fig. 2F). The extra-corporeal system of vessels is an essential part in the colonial strategy in *Symplegma*. Blood vessels systems maintain cohesion and homeostasis of the colony. Vessels have the capacity to rearrange in case of external disturbance (Gutierrez & Brown, 2017)□; or by the dormancy at cold season, by the resorption of all zooids maintaining only the blood vessel system (Hyams et., 2017). The plasticity of the blood vessels system is decisive in the capacity of self-organization of the colonies. As a result *Symplegma* colonies are more resilient to external disturbances, in comparison with solitaries species.

### 4.2. Blood cell types distribution regulates the modularization of developmental processes in colonies

The blood cell types described in *S.rubra* (FIG.3) are very similar to *S.brakenhielmi* and botryllids, as well as other phylogenetically more distant colonial tunicates (Cima et al., 2001; Gutierrez & Brown, 2017; Hirose et al., 2003)□. These results sugges that this variety of blood cells are related with colonial life history.

One of the main characters of *Symplegma* coloniality is the simultaneous budding stages. The blood cells are continuously circulating and their proportions are constant all the time around the colony. In contrast to the botryllids, in which the blood cells proportions have fluctuating cycles in relation with the budding stage of the colony (Ballarin, Menin, et al., 2008)□. In botryllids all the zooids are in the same stage, because the asexual development of the zooids is synchronized (Lauzon et al., 2002)□.

In *Symplegma* double vesicle stage (st.3) hemoblasts (HE) and hyaline amebocytes (HA) were significantly more frequent, than in aging zooids (Fig. 4, Fig 6). In addition, macrophage-like cells (MLC) and morula cells (MC), were significantly more frequent in aging zooids. Suggesting a different location of the blood cells types related with the stage of the zooids and the specific developmental processes occurring in each stage. Thus, precursor cells (HE, HA) are located in buds during the morphogenesis. Probably these cells interact with the double vesicle epithelia in the cellular differentiation and migration before organogenesis (Brown et al., 2009)□. Phagocytes (MLCs) and cells for allorecongition (MCs) are located predominately in the aging zooids. The phagocytes have and active role in the resorption and recycling tissues of the aging zooids (Lauzon et al., 1993).

Dynamic cellular behaviors of amebocytes and morula cells, can be involved in the biological function of these cells (i.e buds morphogenesis and immune responses). These cells migrate in the blood cells and inside zooids and buds, regulation budding, regeneration and immune responses. This faster cellular movements can be a factor related with plasticity of colonies to external disturbances. As well is a interesting source of unexplored biological information.

### 4.3. Aging in Symplegma is a regulate process involving programmed cell death and phagocytosis

The steps described in the stage 11 in *Symplegma* are very similar with the resorption stage described for botryllids (Ballarin, Burighel, et al., 2008). Beginning with the siphons closure, followed by the disintegration of epitheliums and an accumulation of MLCs around the pharynx and stomach (Fig. 5). Posteriorly, the epithelium cells and internal tissues start massive apoptosis. Finally these apoptotic bodies are reabsorb by phagocytes to move them outside from the aging zooids. Thus, the programmed aging of the old zooids and the resorption of these tissues by the colony are processes that evolve in the clade *Symplegma + Botryllids* (Ballarin, Menin, et al., 2008; Lauzon et al.,1992)□. Suggesting the develop of new and complex biological processes related to colonial life history. This programmed aging and the recycling of old zooids by this phagocytosis serie are innovations of colonies, suggesting an increase in the complexity pattern in the evolution of coloniality.

In conclusion this results support the hypothesis that the regulation of the simultaneous developmental processes in *Symplegma* is related by the distribution and modulation of this variety of blood cells types. Therefore, it is probable that the regulation of budding is also related with vascular and zooids epitheliums. This regulation could be mediated by committed epitheliums, which interact with undifferentiated blood cells to start the asexual development.

### 4.4. Evolution of colonial life history in Styelidae: a case of natural selection favoring increase in complexity?

The colonial life history evolved in Styelidae by different developmental mechanisms, during two independent events (Alié et al., 2018)□. However the concept of transform the unique individual in solitaries in a clonal module, it is convergent in the different colonial strategies. This colonial life history evolved by convergence in other marine pelagic animals. Such as, cnidarians, bryozoans and hemichordates (Davidson et al., 2004)□.This support the idea that in some marine environments the coloniality can be a successful strategy.

Specifically in Styelidae the organisms are sessile filter-feeders, thus colonial strategy can give some advantages to survive. The colonies act as self-regulating systems that can rearrange it components (i.e zooids and blood vessels) to maintain the homeostasis in case of a disturbance. An example of this self-organization is the regeneration process. when a portion of the colony is lost or in the whole-body regeneration. In which the remanent modules of the colony ( i.e., zooids, buds and vascular tissues) rearrange themselves to replace the lost parts (Brown et al., 2009; Gutierrez & Brown, 2017).

Evolution of coloniality from a solitary ancestor involved an increase in complexity, such as Styelidae example. Therefore, in colonial animals increase the number of modules, number of biological hierarchies (e.g colonial hierarchy) and nestedness processes (e.g sexual reproduction and budding to form colonies). In Styelidae complexity increases with evolution of colonial animals, also coloniality evolved and probably disappear several times in tunicates (Kocot et al. 2018)□ and in general in metazoans (Davidson et al. 2004; Scrutton 2015; Hiebert et al. 2020)□, these events involving changes in the complexity patterns. Life evolved without a directionality in a spectacular diversity of life forms, the study of complexity patterns can provide a useful way to understand the process of life evolution.

## Acknowledgments

This research was supported by CAPES Brazilian agency, and the grant nº 2015/22650-2 from São Paulo Research Foundation (FAPESP). Many thanks to the *CEBIMar,* Laboratory of Protist evolution, and EvoDevo Laboratory, from Universidade de São Paulo for the technical support.

**Video. 1 Blood cells behavior from Symplegma rubra.** Majority of the cells are morula cells, which a circular shape that change with the movements of the cells. Amebocyte located in the middle, has a dynamic movemnt, at the final of the video pseudopod are visible. Living cells are with natural colors, in the extracellular matrix laminin. Link to watch the video: https://drive.google.com/file/d/14anft05kpxk5HxjXKtTeZ3ihgKDoWlq4/view?usp=sharing.

**Video. 2 Amebocytes behaviors.** Three amebocytes are showing dynamic movements, pseudopods are visible. Cellular movements are related with drasticall change in cytoplasm shape and pseudopod extensions. Living cells are with natural colors, in the extracellular matrix laminin. Link to watch the video: https://drive.google.com/file/d/1HXQSVc6jCwvHBmuAek02bngN11eaJ6m5/view?usp=sharing.

